# Local adaptation shapes community composition without affecting performance

**DOI:** 10.1101/2025.06.06.658317

**Authors:** Swapna Krithika Subramanian, Daniel I Bolnick

## Abstract

Eco-evolutionary theory predicts that local adaptation should influence community assembly and performance, yet empirical tests of these predictions remain limited, particularly at microgeographic scales where gene flow is expected to constrain adaptation. Using yeast communities in apple orchards as a model system, we tested whether local adaptation of a dominant species affects community composition and performance through reciprocal transplant experiments across eight Connecticut orchards and three apple varieties. The dominant yeast *Hanseniaspora uvarum* varied dramatically in abundance (4.3-97.5%) and exhibited significant local adaptation and maladaptation patterns across orchard-variety combinations. Critically, *H. uvarum* local adaptation strongly predicted community composition: locally adapted populations achieved higher abundance while suppressing competitors. However, local adaptation had no effect on community-level performance in apple decomposition. Communities dominated by locally adapted *H. uvarum* performed equivalently to diverse assemblages, indicating functional redundancy among species. These results demonstrate that rapid evolutionary change can decouple community composition from performance, suggesting that functional redundancy can maintain community performance despite dramatic compositional shifts driven by local adaptation.

## Introduction

Local adaptation can reshape species interactions and drive community assembly, particularly in systems where evolution occurs rapidly enough to influence ecological dynamics (Hairston et al. 2005; De Meester et al. 2016; Foster et al. 2017). As species adapt to their local environments, they may alter competitive hierarchies, resource use patterns, and ultimately community composition and performance. These eco-evolutionary dynamics have been documented across a variety of taxa and spatial scales, from old-field plants (Crutsinger et al. 2007), to arthropod communities (Farkas et al. 2013), to freshwater zooplankton (De Meester et al. 2007; Pantel et al. 2015). However, their importance remains largely unexplored at microgeographic scales where high gene flow between nearby populations might be expected to hinder local adaptation (De Meester et al. 2016).

Theory suggests that local adaptation could impact community composition through multiple mechanisms, depending on how species utilize and compete for resources. For instance, early colonizers that become locally adapted might monopolize key resources, leading to community domination by the early-arriving adapted species (De Meester et al. 2002, Urban and De Meester 2009). Alternatively, local adaptation can promote coexistence through niche partitioning (Lawrence et al. 2012), or through shifts in competitive abilities that balance species abundances when resource use between species cannot be partitioned (Hart et al. 2019). These contrasting scenarios suggest that evolutionary dynamics can either increase or decrease community diversity, depending on how adaptation alters species interactions and resource use patterns.

Local adaptation may also influence community-level performance, which we define here as the community’s collective ability to utilize available resources. When local adaptation leads to community monopolization by one species, the loss of functional diversity could decrease performance by potentially losing a species that uses a specialized resource or facilitates more even resource use and reducing the portfolio effect that typically buffers communities against environmental variation (Loreau and de Mazancourt 2013). Conversely, niche partitioning driven by adaptation could enhance productivity through evolving complementary use of resources and cross-feeding (Lawrence et al. 2012). A third possibility is that community composition changes might have no effect on performance if species are functionally redundant in their resource use capabilities (Schindler et al. 2015). These theoretical predictions have rarely been tested empirically in natural systems, particularly at microgeographic scales where rapid evolution might most strongly influence community assembly (Richardson et al. 2014).

Yeast in fungal microbiomes, with their short generation times and capacity for rapid evolution, offer a promising system to study eco-evolutionary dynamics (Lawrence et al. 2012; Nemergut et al. 2013). Fungal microbiomes, also known as mycobiomes, often house a diversity of yeasts. Wild yeasts are also known to have more genetic variation than is present in the many strains propagated in labs (Liti 2015), providing more reason to study their local adaptation in the wild. Despite their ecological and evolutionary significance, fungal communities are often studied primarily for their industrial implications, as certain fungi can ruin crops and alter alcoholic products irrevocably (Shen et al. 2018; Abdelfettah et al. 2021).

Fruits, as sugar-rich environments, are common habitats for fungi, and their microbial communities have played a key role in human domestication of yeast for brewing. There are a variety of studies on yeast community composition in alcoholic fermentations, from traditional Chinese liquor (Wu et al. 2013), tequila (Verdugo Valdez et al. 2011; Lopez et al. 2014), cocoa beans (Mota-Gutierrez et al. 2018) and the most studied, grape wines (Nisiotou et al. 2007; Gayevskiy and Goddard 2012; Milanovic et al. 2013). Therefore, a large portion of yeast community studies focus on the interaction of introduced *Saccharomyces cerevisiae* strains with the wild yeast already present on the fruit before fermentation. Gayevskiy and Goddard (2012) studied yeast communities on grapes before and after fermentation in New Zealand and found regional differences in yeast communities. Milanovic et al. (2013) also studied yeast communities on grape surfaces before fermentation and after fermentation and additionally found that vineyards that sprayed fungicide had lower yeast diversity than organic vineyards.

Efforts to characterize microbiomes in apples, show that apple microbiomes vary widely by country, orchard, and even fruit tissue type (Abdelfettah et al. 2021). Additionally, Abdelfettah et al. (2016) characterized the fungal community of Red Delicious apples in supermarkets, finding significant differences between fruit parts and between organic and conventional apples, though no significant effect of two weeks of storage. Shen et al. (2018) specifically studied fungal communities on apples before and after cold storage, showing that changes in storage conditions could cause postharvest deterioration. While apple fungal communities have been studied in the context of production, the effect of local adaptation on fungal community composition and performance has remained unevaluated.

Apple orchards are a useful model system for studying the impacts of local adaptation on microbial communities. In the northeastern US, many apple orchards have been established and growing apples for eight or more generations of farmers, spanning hundreds of years of careful records and practices. This long-term operation has given wild yeast populations ample time to evolve in response to orchard and variety-specific conditions. Apples create small, sugary environments that airborne yeast can colonize through wind dispersal or insect dispersal and can subsequently rapidly adapt to. These apples form a series of replicated habitats spanning a range of increasing spatial scales: many apples within a tree, many trees within a particular grove, multiple apple varieties within an orchard, and multiple orchards within a geographic region. Yeast might adapt to local conditions at any of these scales, but particularly interesting is whether this could result in adaptation at the microgeographic scales at which you would assume there would be genetic mixing due to their large dispersal neighborhood, such as the scale of varieties within an orchard (Richardson et al. 2014). Orchard-specific traits, like the local climate and soil are a major source of environmental variability. Orchards also often employ different types and frequencies of fungicide spraying, and research has shown that fungicide use in vineyards can lead to reduced yeast diversity (Milanovic et al. 2013), while untreated orchards retain a greater diversity of wild yeasts. Apple varieties introduce another source of variation and can differ in several traits affecting yeast performance within and across species, including sugar and amino acid profiles, fruit firmness, and cellular structure (Lapsley et al. 1992; Wu et al. 2007). Due to the clonal nature of apple varieties, this variation is replicated across orchards, potentially resulting in parallel adaptation despite environmental differences. With farmers providing selection pressures and the apple varieties providing replicate environments, apple orchards can be considered a long-term semi-natural evolution experiment for yeast. This microgeographic adaptation could play a significant role in the composition and performance of fungal communities within fruits.

This study tests whether 1) yeast communities in apple orchards differ between orchards and varieties, and 2) whether these communities perform better on their native orchards/varieties via reciprocal transplant assay. We identify a dominant community member within the characterized yeast communities and use a reciprocal transplant assay to 3) determine if the selected yeast species is locally adapted to different orchards and varieties. Finally, we 4) test whether the variation in community composition and performance is explained by the local adaptation of the selected dominant species.

## Methods

### 1) Do yeast communities vary among orchards and varieties?

We characterized the apple fungal community in rotting apples from eight different orchards in Connecticut (USA) (Supplemental Table 1) and 3 different varieties within those orchards. We first collected rotting apple and fresh apple pairs from the same tree for every orchard-variety combination. We handled apples using sterile techniques in the orchards before taking them back to the lab. We selected rotting apples by evaluating whether they had an opening in the flesh that was caused by natural puncturing and that the puncture was facing up rather than in the soil to avoid extraneous soil microorganisms. We selected fresh apples by ensuring that they were attached firmly by the stem to the tree and that there were no punctures in the skin, and we sterilized the skin with ethanol before taking them back to the lab. For each orchard-variety combination, we collected five replicates (rotting and sterilized fresh apples), all from different trees.

In the lab, we flame-sterilized a metal inoculating loop and swirled it in the rotting flesh within the punctured rotting apples and streaked the yeast community on a YPD agar plate with 1% penicillin-streptomycin to suppress bacterial growth. After allowing the plate to culture for 24 hours, we randomly chose eight isolated colonies using 20 uL sterile pipette tips to culture in YPD media. These eight isolates represented a random sub-sample of the fungal community for each orchard-variety combination in each replicate. We cultured the yeast isolates for 24 hours and then conducted a BOMB DNA extraction (Oberacker et al. 2019). To identify each yeast isolate, we conducted ITS region barcoding using ITS1 and ITS4f primers and send the PCR products for Sanger sequencing (Martin and Rygiewicz 2005).

We calculated Simpson’s diversity index for each replicate rotting apple community using the “vegan” package (Okansen et al. 2022) in R (v4.4.1, R Core Team 2024). While we acknowledge that our sampling of eight isolates per apple represents a limited sample for diversity estimation, we chose Simpson’s diversity index because it is more robust to sampling limitations, particularly for communities dominated by a few species (Gotelli and Colwell 2001; Roswell et al. 2021). We tested whether Simpson’s diversity varied among orchards and varieties using a linear model with orchard, variety, and their interaction as fixed effects (Table 1a). Type III sums of squares ANOVA was performed on the linear model and results were interpreted to identify significant differences between orchards and varieties. The impact of orchard and variety differences on community composition was quantified using redundancy analysis (RDA) in the R package “vegan” with orchard, variety, and their interaction as fixed effects. Community data was transformed using Hellinger transformation to account for species abundance and the model was fitted. Permutation-based ANOVA with 999 permutations was used to test for significance of terms, and total variance (R^2^) explained by the model was quantified. Using the above analyses, we identified *H. uvarum* as the most abundant and most variable species for further study of local adaptation effects.

**Table 1:** Community composition statistical model results. a) Simpson’s diversity index (α) for yeast communities across orchard and variety combinations, b) community performance evaluated by rot size, home vs away (by yeast source) Type III ANOVA results on linear models focused on orchard and variety, c) community performance evaluated by rot size, local vs foreign (by apple source) Type III ANOVA results on linear models focused on orchard and variety.

| a) Simpson's diversity index ($\alpha$) | | | |
| --- | --- | --- | --- |
| Factor | df | F value | p value |
| Orchard | 7,92 | 4.33 | <0.01 |
| Variety | 2,92 | 1.09 | 0.34 |
| Orchard*Variety | 13,92 | 1.61 | 0.10 |
| b) community performance, home vs away (by yeast source) |  |  |  |
| Factor | df | F value | p value |
| Native/Non-native Orchard | 1,2376 | 3.15 | 0.08 |
| Yeast Source-orchard | 7,2376 | 10.33 | <0.01 |
| Native/Non-native Orchard*<br>Source-orchard | 7,2376 | 6.55 | <0.01 |
| Native/Non-native Variety | 1,2386 | 0.49 | 0.49 |
| Yeast Source-variety | 2,2386 | 8.82 | <0.01 |
| Native/Non-native Variety*<br>Source-variety | 2,2386 | 4.00 | 0.02 |
| c) community performance, local vs foreign (by apple source) |  |  |  |
| Factor | df | F value | p value |
| Native/Non-native Orchard | 1,2376 | 4.90 | 0.03 |
| Apple Source-orchard | 7,2376 | 10.88 | <0.01 |
| Native/Non-native Orchard*<br>Source-orchard | 7,2376 | 3.86 | <0.01 |
| Native/Non-native Variety | 1,2386 | 4.49 | 0.03 |
| Apple Source-variety | 2,2386 | 8.83 | <0.01 |
| Native/Non-native Variety*<br>Source-variety | 2,2386 | 4.40 | 0.01 |

### 2) Does yeast community performance differ among orchards or varieties?

We conducted a reciprocal transplant experiment with the yeast communities (sampled from rotting apples) to test their performance within the apples (Fig. 1a). We define ‘performance’ as the extent of apple tissue degradation caused by yeast activity, measured by the spread of visible browning (Kidd and Beaumont 1924, Bano et al. 2023). As yeasts consume sugars and spread through the apple tissue, they break down cell walls through fermentation byproducts such as ethanol and acids. This cellular damage leads to oxidation and tissue browning and allows yeast to spread to neighboring cells and consume more sugars. Therefore, the area of brown tissue indicates how successfully the yeasts have colonized and metabolized the apple substrate, with larger areas of browning indicating higher yeast performance. To create a representative community of each orchard-variety combination, we pooled equal concentrations of each yeast isolate within each orchard-variety combination i.e. eight isolates across five replicates, or 40 isolates, and reciprocally transplanted them into the fresh apples. To pool the yeast in equal concentrations, we cultured the isolates for 24 hours and then measured OD600. This culturing period in addition to previous periods where the yeast grew in the lab act as a common garden period to avoid inherited environmental effects. We then pooled each isolate in a concentration adjusted for their OD600 so each community has an equal concentration of each yeast isolate and across communities.

**Figure 1:**
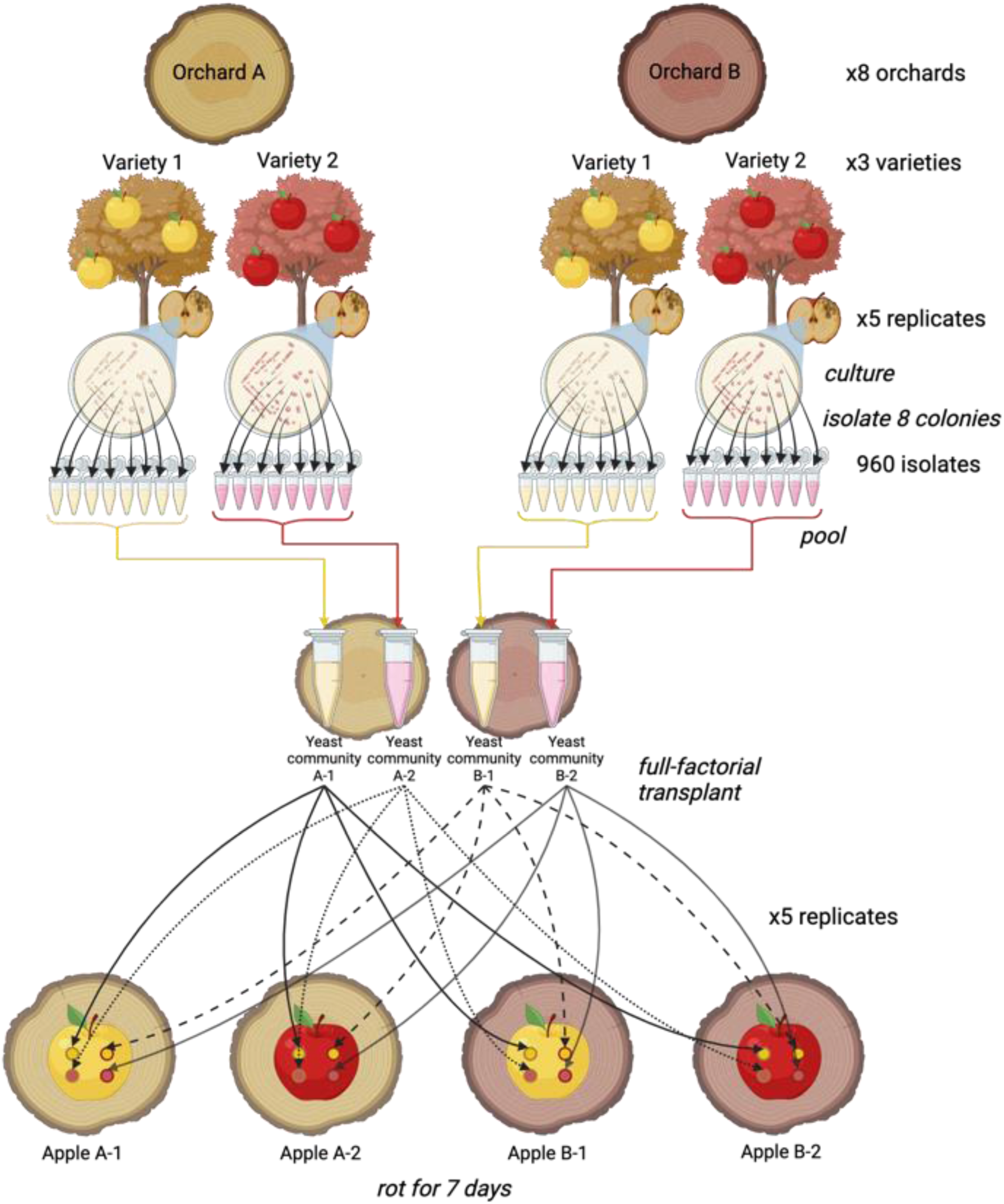
Reciprocal Transplant Assay Experimental Methods. Community performance experimental methods. Created using Biorender.

To inoculate the apples with yeast communities, we first sterilized the skin of the fresh apples using ethanol. Using a sterile 200 uL filter pipette tip, we stabbed the apple to a depth of 21 mm (the 100 uL mark on the tip) and then inoculated the resulting hole with 50 uL of the respective yeast community. We inoculated five replicates of fresh apples of all orchard-variety combinations with yeast communities of all orchard-variety combinations (Fig. 1a). After inoculation, we sealed the hole with microporous sealing film to allow for aeration and to prevent extraneous yeast from contaminating the communities and to allow the yeast to conduct aerobic respiration from within the apple as they would in the orchards. Each apple additionally had a control hole that did not receive any yeast, to serve as an indicator for if the apple is contaminated by extraneous yeast. Contamination did not occur on any apples. After 7 days, we measured performance as the longest diameter of the brown flesh (rot) that developed beneath the apple skin around each inoculation hole using digital calipers in inches.

For community performance analyses, we used separate linear models for orchards and varieties to avoid collinearity issues. To analyze performance in relation to orchards, we used rot size as the dependent variable with native/non-native orchard transplant, yeast’s source-orchard, and their interaction as fixed effects. Similarly, for variety analyses, we used rot size as the dependent variable with native/non-native variety transplant, yeast’s source-variety, and their interaction as fixed effects. These models tested local adaptation using the “home versus away” criterion (Kawecki and Ebert 2004), which compares performance in native versus non-native environments, focusing on a constant yeast community. We also ran complementary models using the “local versus foreign” criterion (Kawecki and Ebert 2004) by including the apple’s source instead of the yeast’s source as a predictor. To explore relationships between community diversity and performance, we additionally ran a linear model with rot size as the dependent variable and Simpson’s diversity index, orchard, and variety as fixed effects. We conducted ANOVAs with Type III sums of squares on all models and calculated standardized effect sizes using the “lm.beta” package (PedersenBehrendt 2014).

### 3) Does a dominant community member (H. uvarum) exhibit local adaptation?

We used the ITS region barcoding results from the collected and characterized yeast communities to identify *H. uvarum* as a common yeast in the apple orchard community. We then selected five isolates, each from a different replicate apple across the eight orchard-variety combinations, i.e. 40 *H. uvarum* strains to test for local adaptation (Fig. 1b). If a rotting apple replicate community did not have an *H. uvarum* strain, a replacement strain was randomly selected from all *H. uvarum* strains in that orchard-variety combination. We conducted a reciprocal transplant experiment with the five isolates from eight orchard-variety combinations. We cultured each of the 40 isolates in YPD media for 24 hours and then quantified their OD600. We then diluted each isolate with fresh YPD so each isolate will start at equal concentration in the apples. We sterilized the skin of the fresh apples using ethanol. Using a sterile 200 uL filter pipette tip, we stabbed the apple to a depth of 21 mm (the 100 uL mark on the tip) and then inoculated the resulting hole with 50 uL of the respective yeast isolate. We inoculated five replicates of fresh apples of all orchard-variety combinations with five isolates of all eight orchard-variety combinations replicated three times each (Fig. 1b). After inoculation, the hole was sealed with microporous sealing film. Each apple additionally had a control hole that did not receive any yeast. *H. uvarum* isolate performance was measured using digital calipers in inches after seven days as the longest diameter of the brown flesh (rot) that developed beneath the apple skin around each inoculation.

For *H. uvarum* local adaptation analyses, we used a similar approach to the community performance analyses but ran a mixed effects model including yeast strain as a random effect. This approach was chosen because strains were randomly sampled from each orchard-variety combination and represent a broader population of potential strains. The orchard analysis used rot size as the dependent variable with native/non-native orchard transplant, yeast’s source-orchard, and their interaction as fixed effects, and individual yeast strain as a random effect. The variety analysis similarly used rot size as the dependent variable with native/non-native variety transplant, yeast’s source-variety, and their interaction as fixed effects, and individual yeast strain as a random effect. As with the community analyses, we tested both “home versus away” and “local versus foreign” criteria (Kawecki and Ebert 2004) by running parallel models with either yeast source or apple source as predictors. We fit linear mixed-effects models using the “lmerTest” package (Kuznetsova et al. 2017), which provides p-values for fixed effects using Satterthwaite’s method for denominator degrees of freedom. We assessed significance using Type III analysis of variance and calculated standardized effect sizes and confidence intervals using the “effectsize” package (Ben-Shachar et al. 2020).

### 4) Does H. uvarum local adaptation explain variation in community composition or performance?

Log fold change (LFC) in rot size between native inoculations and non-native inoculations was calculated for all *H. uvarum* strains as a measure of local adaptation, and for all communities as a measurement of performance. It was calculated with native values/non-native values. The influence of *H. uvarum* local adaptation on community performance was determined by running a linear model correlating LFC of communities to LFC of *H. uvarum* strains. A positive relationship would be consistent with *H. uvarum* abundance in communities increasing as they are more locally adapted. *H. uvarum* local adaptation influence on community composition was evaluated comparing RDA models with and without *H. uvarum* strain LFC as a fixed effect. RDA models were compared using total variance explained (R^2^) and p-values from permutation-based ANOVA with 999 permutations.

## Results

### 1) Yeast community composition varies across orchards and varieties

Yeast communities varied systematically across different orchards and varieties, with diversity and composition showing distinct spatial patterns. Among the species identified, *Hanseniaspora uvarum* emerged as the most abundant species, though its prevalence varied dramatically across sites, from 4.3% of the community in Orchard H Empire apples to 97.5% in Orchard B Golden Delicious apples (Fig. 2). Other frequently occurring species included two *Pichia* species (*P. fermentans* and *P. kluyveri*) and *Metschnikowia pulcherrima*, but were not as abundant as *H. uvarum*. Overall yeast abundance across all samples can be found in the apple pie chart in the Supplementary Materials (Supplemental Figure 2)

**Figure 2:**
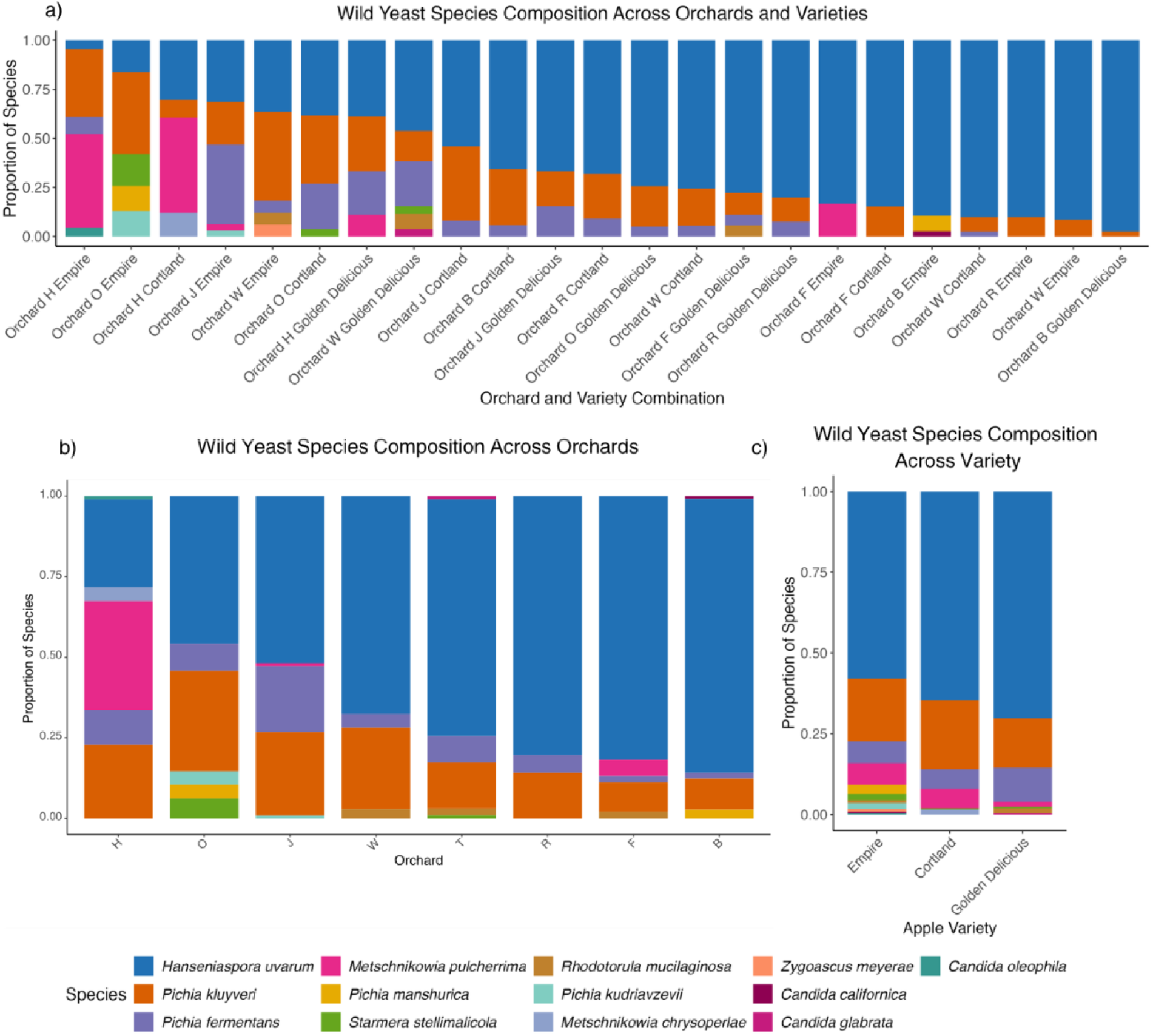
Species Composition of Yeast Communities Across Orchards and Varieties. Relative abundance of yeast species across eight orchards and three apple varieties. Bars show the proportion of each species within community samples.

Community diversity, measured by Simpson’s index, varied significantly among orchards but not among varieties or their interaction (Table 1a). Diversity varied 15-fold from 0.049 in Orchard B Cortland apples to 0.74 in Orchard O Empire apples (Table 1a, Fig. 3).

**Figure 3:**
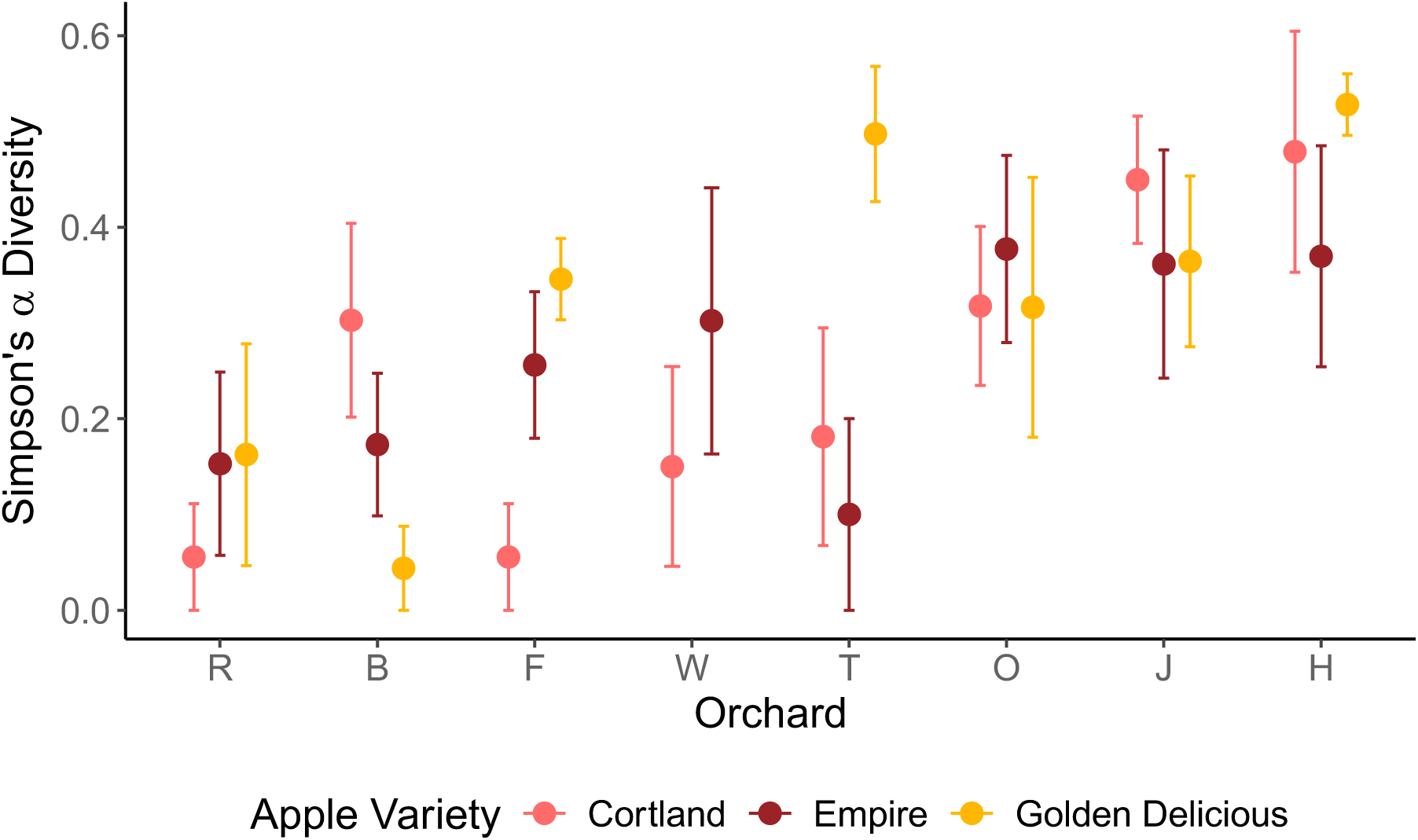
Simpson’s Diversity Index Varies Among Orchards. Simpson’s diversity index (α) for yeast communities across orchard and variety combinations.

Redundancy analysis revealed that community composition was significantly structured by both source-orchard and orchard-variety interactions. The RDA model explained 39.6% of total community variation, with the first two axes capturing 23% and 9.5% of constrained variance respectively (Fig. 4). The RDA model (*F_22,92_*=2.75, *p*=0.001) and the first RDA axis were significant (*F_1,102_*=38.44, *p*=0.001), while the second RDA axis was not significant (*F_1,102_*=15.97, *p*=0.11). Orchard effects strongly influenced community structure (*F_7,92_*=4.44, *p*=0.001), and while variety alone showed marginal effects (*F_2,92_* =1.99, *p*=0.07), the orchard-variety interaction was significant (*F_13,92_*=1.95, *p*=0.002).

**Figure 4:**
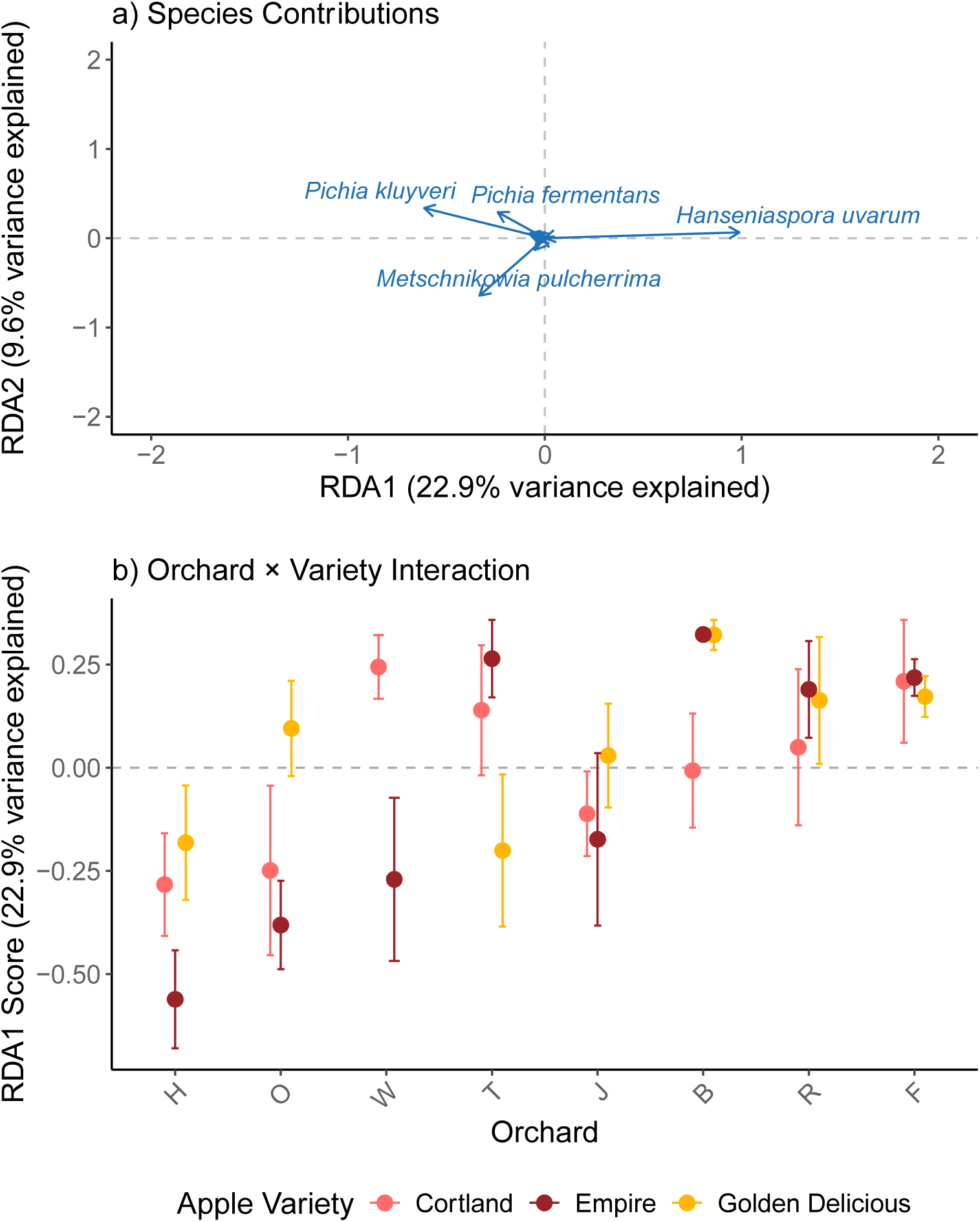
Redundancy Analysis (RDA) of Yeast Community Composition. a) Species-environment biplot showing relationships between individual species abundances (blue arrows), where the length and direction of arrows indicate the strength and direction of species’ contributions to community variation. b) RDA1 scores by orchard-variety combinations demonstrating how community composition varies across different environmental conditions. RDA1 and RDA2 together explain 32.5% of total community variation.

The occurrence of *H. uvarum* is negatively correlated with the occurrence of the other most abundant species and seems to be a strong driver of the variation across the RDA1 axis (Fig. 4a). While many of the species across the communities lay around zero in the RDA plot, the focal species *H. uvarum* had the most extreme position on the significant axis RDA1, with a score of 0.99 (Fig. 4a), while the two next most abundant species in the *Pichia* genus were both on the negative end of the RDA1 axis. The orchard RDA1 scores are consistent with the abundance of *H. uvarum*, with orchards where *H. uvarum* are common having positive scores (Orchard B, Orchard F, Orchard R) and orchards where *H. uvarum* is rare having negative scores (Orchard H, Orchard O, Orchard J, Orchard W). The interaction of orchard with variety was complex, with varieties having different rank order in orchards (Fig. 4b). For example, while in Orchard H the Empire apples had the lowest *H. uvarum* abundance and the Golden Delicious had the highest *H. uvarum* abundance, the pattern was exactly the opposite in Orchard T, with the Golden Delicious having the lowest *H. uvarum* abundance and the Empires the highest (Fig 2, 4b).

### 2) Yeast community performance varies across orchards and varieties

Yeast community performance varied by orchard and variety, with some doing better at decomposing apple tissues from native orchards/varieties and some performing better in non-native orchards/varieties (Table 1b, Fig. 5). Yeast source-orchard significantly influenced rot size (Table 1b), with yeasts from Orchard J producing the largest rot sizes overall (β = 0.303, *p* < 0.001, n = 312), followed by yeasts from Orchard B (β = 0.217, *p* = 0.003, n = 312) and Orchard O (β = 0.188, *p* = 0.01, n = 312). Yeasts from Orchard R produced significantly smaller rot sizes overall (β = −0.218, *p* = 0.002, n = 312) (Fig. 5a). While the native vs non-native orchard test was not significant, there was a significant interaction between yeast source-orchard and native vs non-native orchard (Table 1b). Many yeast communities performed better in their native orchards, with Orchard O showing the strongest native advantage (β = −0.239, *p* = 0.001, n = 312), then Orchard J (β = −0.202, *p* < 0.001, n = 312), and then Orchard W (β = −0.157, *p* = 0.04, n = 208) (Fig. 4a). Orchard R was the only yeast community that produced significantly larger rot sizes in non-native orchards (β = 0.181, *p* = 0.01, n = 312) (Fig. 5a). Orchard H and Orchard F showed no significant differences between native and non-native orchards (Fig. 5a).

**Figure 5:**
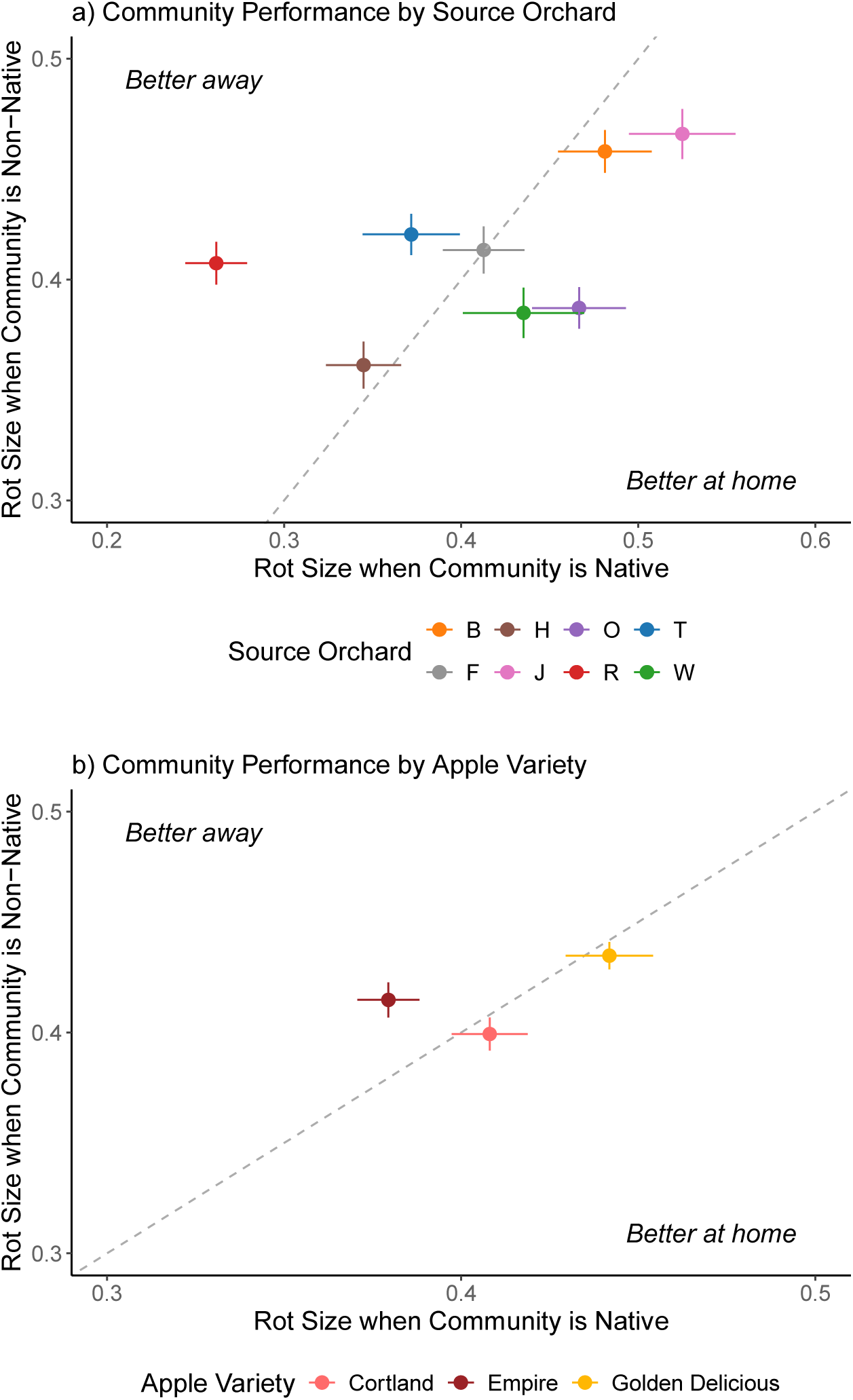
Community Performance in Native versus Non-native Environments. Rot size produced by yeast communities in reciprocal transplant experiments. a) Performance by orchard source. b) Performance by variety source. Error bars represent standard error.

Yeast community performance varied by apple variety, with some doing better in native varieties and some performing better in non-native varieties (Table 1a, Fig. 5b). Yeast source-variety significantly influenced rot size (Table 1a), with yeasts from Golden Delicious generating larger rot sizes overall (β = 0.091, *p* = 0.026, n = 728) and yeasts from Empire showing smaller rot sizes (β = −0.080, *p* = 0.044, n = 832) (Fig. 5b). While native vs non-native variety was not significant, there was a significant interaction between yeast source-variety and native vs non-native variety (Table 1a). Yeasts from Empire apples showed significantly different performance, producing larger rot sizes in non-native varieties (β = 0.108, *p* = 0.012, n = 832), while Golden Delicious performed consistently regardless of whether the variety was native or non-native (β = 0.004, *p* = 0.927, n = 728) (Fig. 5b). Local vs foreign comparisons confirmed that the effects seen are not due to differences in habitat quality (Table 1b). We additionally found no significant correlation between Simpson’s diversity and overall community performance (p = 0.78).

### 3) H. uvarum shows local (mal)adaptation to orchards and varieties

*H. uvarum* performance varied significantly by orchard and whether strains were in native or non-native orchards (Table 2a, Fig. 6a). Accounting for strain identity as a random effect, yeast source-orchard significantly influenced rot size (χ² = 28.56, *p* < 0.001), with Orchard B strains producing larger rot sizes overall (std.coef = 0.66, 95% CI [0.29, 1.04], *p* < 0.001) and Orchard O strains producing somewhat smaller rot sizes (std.coef = −0.35, 95% CI [-0.73, 0.03], *p* = 0.071). While native vs non-native orchard was not significant alone (χ² = 1.51, *p* = 0.22), there was a significant interaction between yeast source-orchard and native vs non-native orchard (χ² = 23.03, *p* < 0.001). Notably, Orchard O strains showed significant maladaptation, producing larger rot sizes when consuming apples from non-native orchards (std.coef = 0.71, 95% CI [0.31, 1.11], *p* < 0.001). Orchard R and Orchard B strains showed no significant differences between native and non-native orchards.

**Figure 6:**
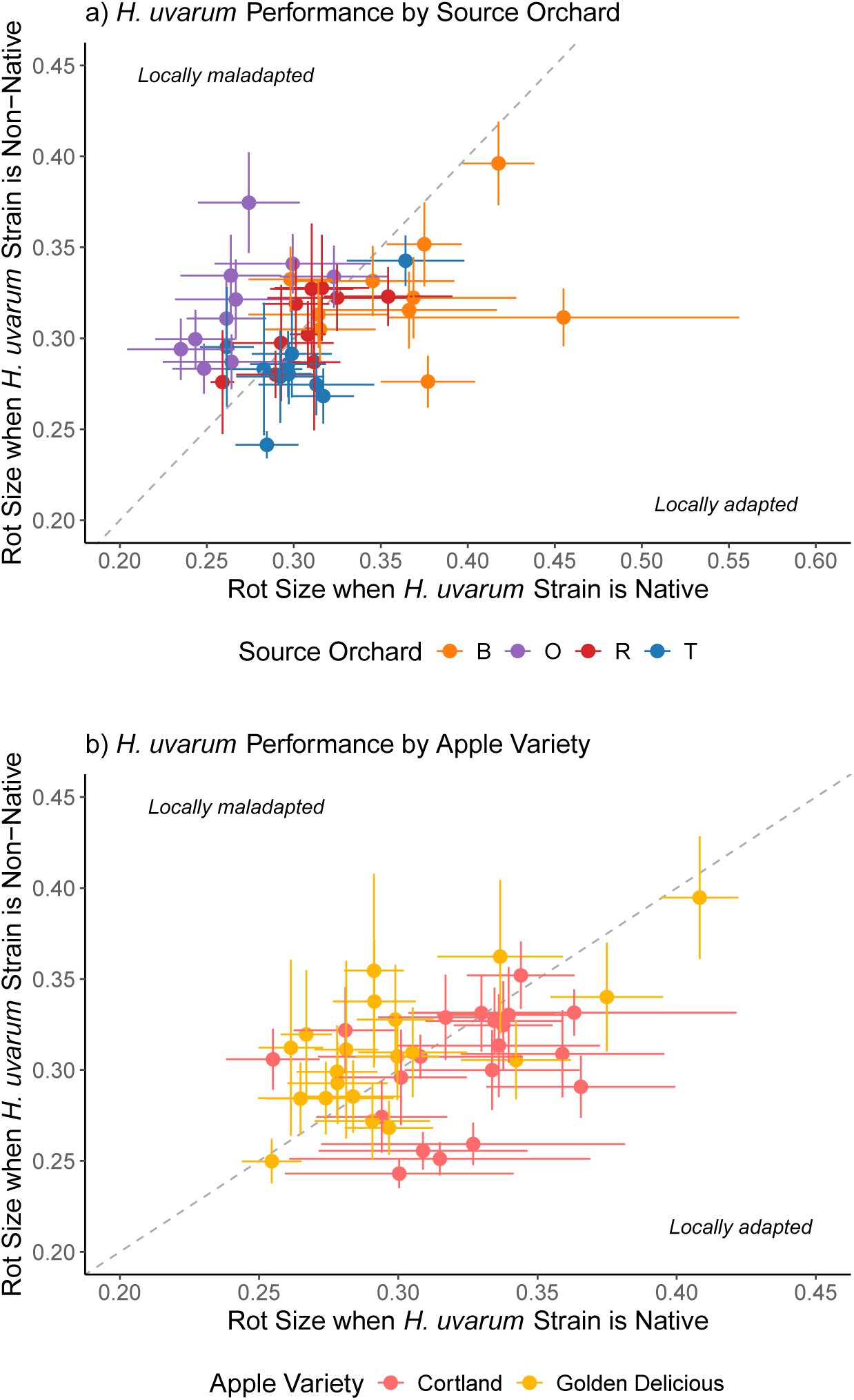
*H. uvarum* Strain Performance in Reciprocal Transplants. Performance of individual *H. uvarum* strains in reciprocal transplant experiments. a) Strain performance across orchards. b) Strain performance across varieties. Error bars showing standard error.

**Table 2:**
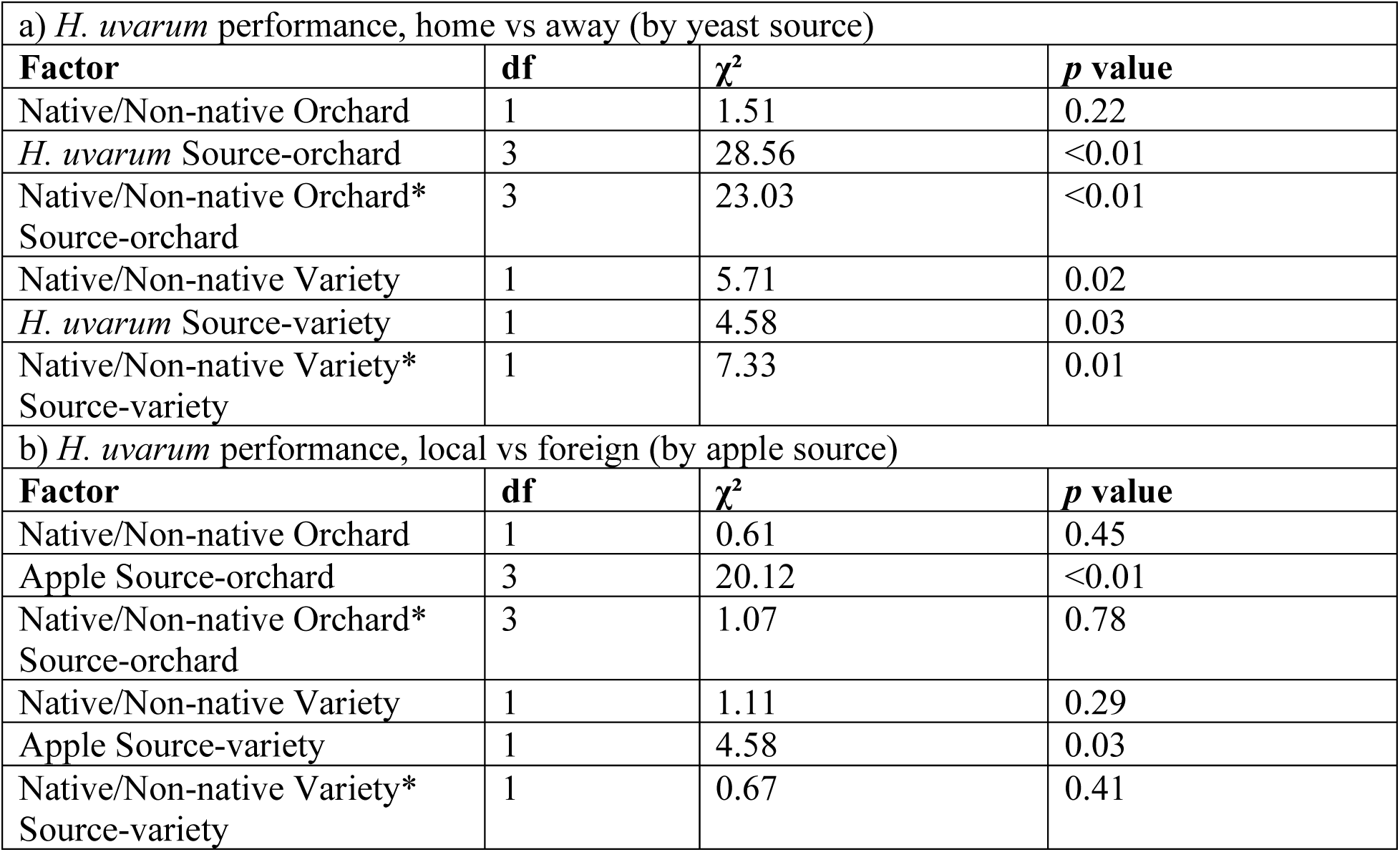
*H. uvarum* local adaptation statistical model results. a) *H. uvarum* performance by Orchard and Variety, home vs away (by yeast source), b) *H. uvarum* performance by Orchard and Variety, local vs foreign (by apple source)

*H. uvarum* performance also varied by the interaction of yeast source-variety and whether strains were in native or non-native varieties (χ² = 7.33, p = 0.01, Table 2a, Fig. 6b). Yeast isolated from Golden Delicious apples demonstrated maladaptation, producing significantly larger rot sizes in non-native varieties (std.coef = 0.34, 95% CI [0.09, 0.58], *p* = 0.01). Because of the periods of growth in the lab, we can be confident these effects are due to heritable genetic differences rather than maternal effects. Local vs foreign comparisons confirmed that the effects seen are not due to differences in habitat quality (Table 2b).

### 4) H. uvarum adaptation shapes community composition but not community performance

The results above establish that yeast communities vary in their performance at decomposing apples, depending on the orchard and variety they are from and tested upon. We also established that *H. uvarum* varies in its performance on apples across orchards and varieties. Given that *H. uvarum* is the dominant member of most of our yeast communities, we hypothesized that the differentiation in community performance is a result of the local adaptation/maladaptation of *H. uvarum* populations within those communities. However, our results lead us to reject that hypothesis: despite *H. uvarum*’s strong local (mal)adaptation patterns and yeast communities’ differential performance in native/non-native transplants by orchard and variety, *H. uvarum* local adaptation did not predict overall community performance (Fig. 7). The linear model correlating community performance and *H. uvarum* LFC in native versus non-native transplants showed no correlation (ANOVA, *F_1,34_*=0.12, *p*=0.75).

**Figure 7:**
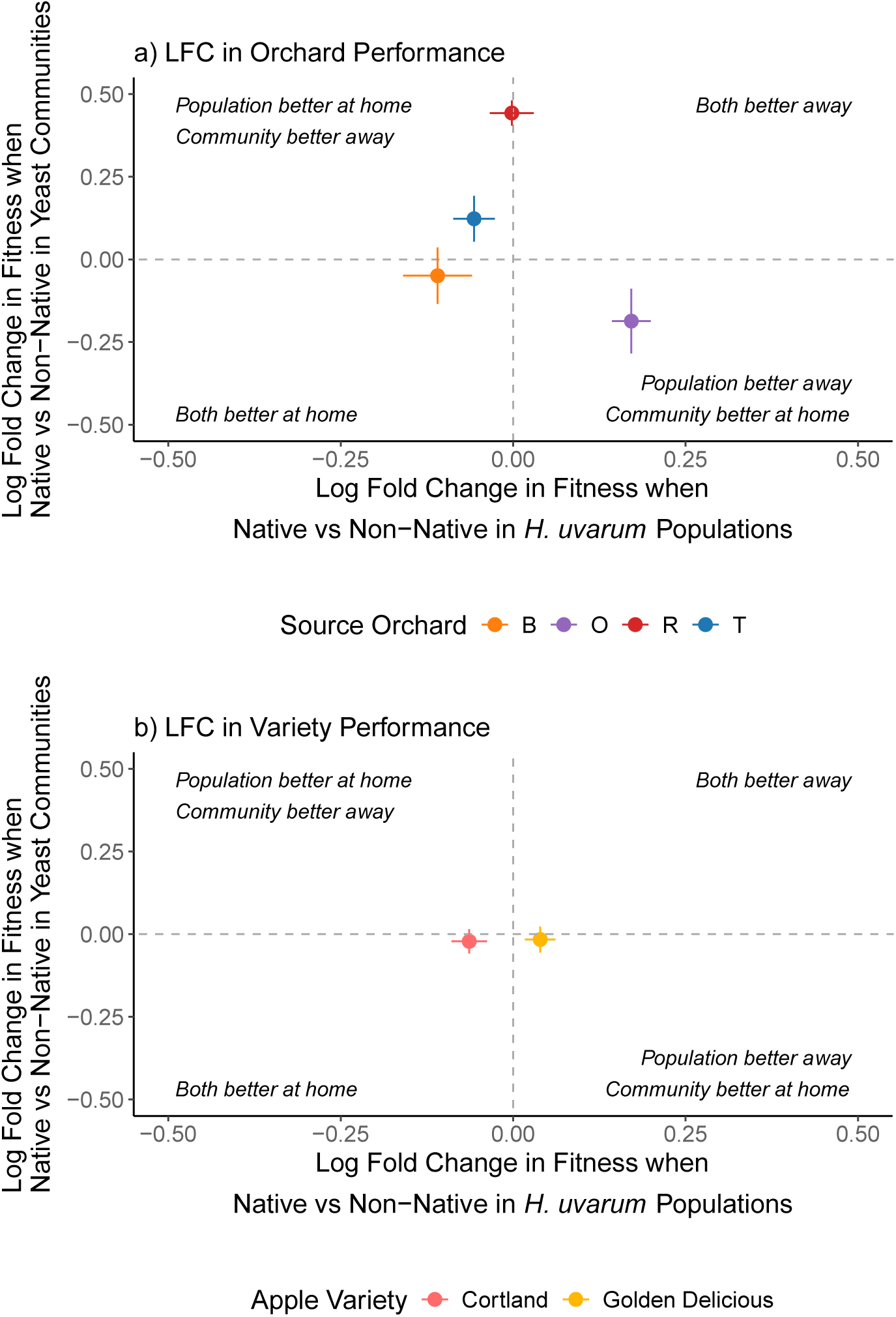
Relationship Between *H. uvarum* Local Adaptation and Community Performance. Log fold change (LFC) in rot size comparing community performance versus *H. uvarum* population performance in a) orchard transplants and b) variety transplants. This figure is showing no significant correlation between *H. uvarum* adaptation and overall community performance.

However, *H. uvarum*’s local adaptation, measured as log fold change in rotting in native vs non-native environments, strongly influenced which species co-occur in the community. Adding *H. uvarum*’s local adaptation patterns to our community composition RDA model dramatically improved our ability to explain community structure, increasing total explained variance from 23.6% (*F_7,28_* = 1.24, *p* = 0.269) to 34.9% (*F_8,27_* = 1.81, *p* = 0.05), and improving model fit (permutation-based ANOVA, *F_1,27_* = 4.66, *p* = 0.01). This improved model showed that *H. uvarum* local adaptation does predict community composition: *H. uvarum* local adaptation is associated with higher *H. uvarum* abundance (RDA1, 27.9% variance, *F_1,30_* = 12.86, *p* = 0.05; *H. uvarum LFC:* 0.524, *H. uvarum*: 0.665; Fig 8a). *H. uvarum* is also associated with lower abundance of the next most abundant species in the dataset, the *Pichia* species (RDA1, *P. kluyveri*: −0.573, *P. fermentans*: −0.394, Fig 8a). When *H. uvarum* is locally adapted to the apple environment, it is better able to outcompete *Pichia* species.

**Figure 8:**
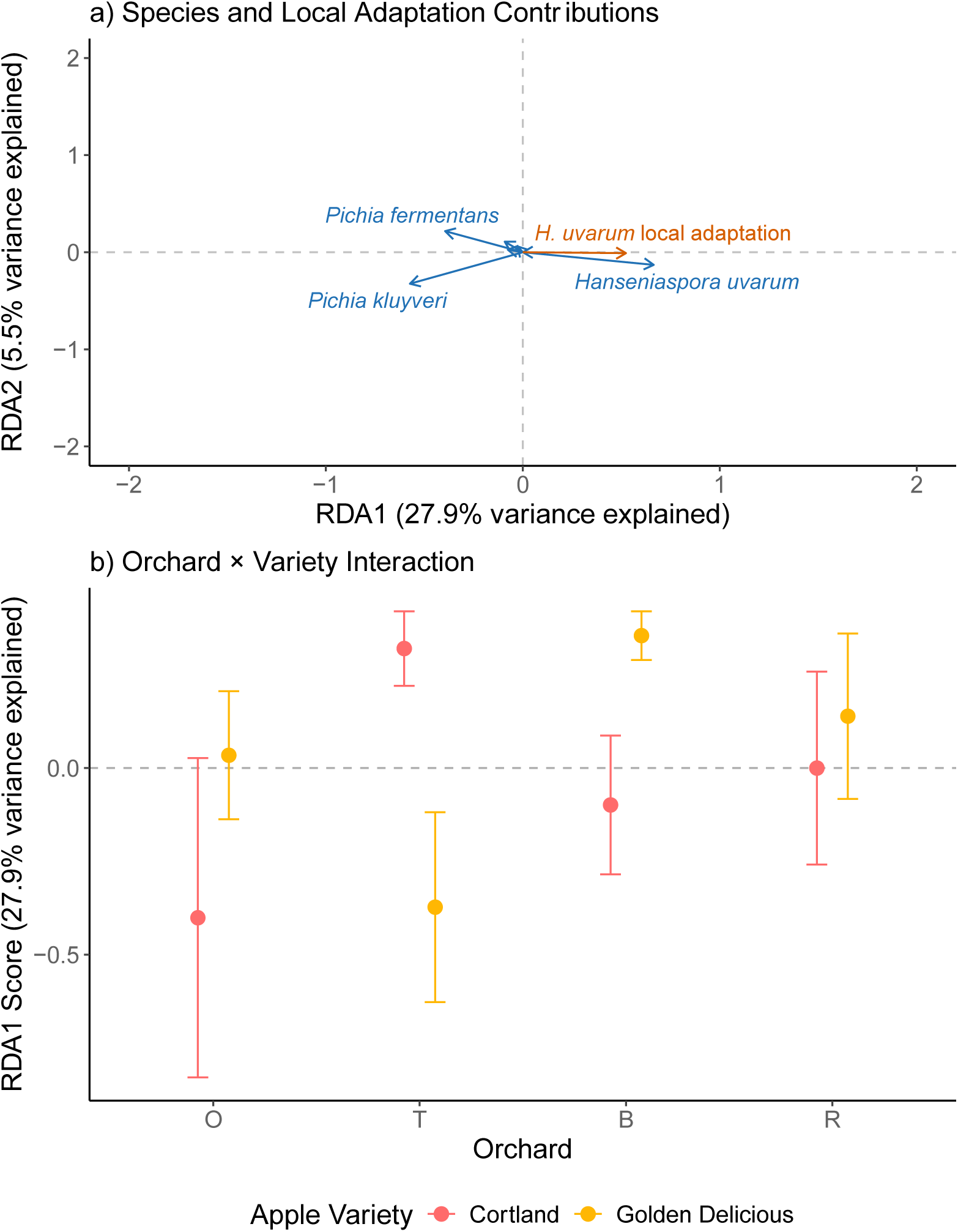
*H. uvarum* Local Adaptation Effects on Community Composition. RDA analysis incorporating *H. uvarum* local adaptation. a) Biplot showing species relationships and how they correlate with *H. uvarum* LFC, with *H. uvarum* showing positive correlation and *Pichia spp.* showing negative correlation. b) RDA1 scores by orchard-variety combinations. The model including *H. uvarum* LFC explains 34.9% of total variance.

Orchard-variety interactions also influenced community composition in the model including local adaptation (*F_3,27_* = 2.96, *p* = 0.02), whereas orchard and variety alone did not (orchard: *F_3,27_* = 0.66, *p* = 0.68; variety: *F_1,27_* = 0.29, *p* = 0.76). For example, Orchard B Golden Delicious apples have the highest positive RDA1 score (0.401) out of all orchard variety combinations, and this is the community in which we observed the highest *H. uvarum* abundance coupled with the lowest competitor abundance (Fig 2, Fig. 8b). These results suggest that while *H. uvarum*’s adaptation doesn’t affect how the communities rot apples, it does reshape community composition, potentially through competitive exclusion. Additionally, the strength and direction of this effect varies depending on the specific orchard-variety combination, and whether the local *H. uvarum* genotypes are well adapted to that combination.

## Discussion

This study explores the effect of local adaptation of an abundant species on community composition and community performance across replicated orchard habitats. Key findings are that 1) yeast communities vary in composition among apple orchards and apple varieties, and 2) their performance in a reciprocal transplant assay also varies across orchards and varieties, with some performing better on home apples and some performing better away. Additionally, 3) the most abundant species across all communities (*H. uvarum*) exhibits local adaptation and local maladaptation to varying degrees across different orchards and varieties. Finally, we find that 4) *H. uvarum* local adaptation significantly impacts community composition without affecting overall community performance. Specifically, *H.uvarum* is more abundant (and competitors like *Pichia spp.* less abundant) in orchard-variety settings where *H.uvarum* is more locally adapted to decomposing the apple resource (in the absence of other yeasts). These findings demonstrate how local adaptation (and maladaptation) can influence community assembly even at microgeographic spatial scales.

### Community Variation and Interactions

The significant orchard x variety interactions in both community composition and performance demonstrate that complex environmental factors are shaping microbial communities at microgeographic scales. For example, while *H. uvarum* dominated the community on Golden Delicious apples from Orchard B (97.5%), it was extremely rare in Empire apples from Orchard H (4.3%). These contrasting patterns reflect the effect of multiple interacting factors. First, differences in orchard management practices, particularly fungicide application regimes, likely create distinct selective environments. While all the orchards tested in this study followed integrated pest management, they each have unique (and proprietary) spraying techniques. Previous work in vineyards demonstrated that fungicide application reduces yeast diversity (Milanovic et al. 2013), which could explain the significant differences in Simpson’s alpha diversity by orchard, but without spraying records we cannot confirm this. Additionally, while all the orchards are located in Connecticut, there could be environmental differences between them that influence the yeast within them. However, orchard differences alone do not explain the complex patterns we observed involving orchard by variety interaction effects. Variety-specific responses suggest that apple chemical composition (Lapsley et al. 1992; Wu et al. 2007) creates distinctive selective environments even within orchards.

The interaction between orchard practices and apple variety characteristics creates a mosaic of selective pressures. For instance, while apple varieties are clonal, the differences in soil nutrients and fertilizer application can alter the chemical composition of apple varieties within the orchards, which might favor different yeast species. The fact that variety effects on community composition varied among orchards suggests these characteristics interact with management practices and environmental factors in complex ways. This aligns with previous work by Abdelfettah et al. (2021) that showed apple microbiomes vary by country, orchard, and tissue type, but reveals that apple variety can add another scale of variation that has not been previously studied.

The spatial structure of these interactions creates opportunities for both local adaptation and meta-community dynamics. While preliminary results showed no significant effects of tree rows (e.g., trees grown in a line, parallel to other rows of the same or different variety; Supplemental Figure 1, Supplemental Table 2), the strong orchard x variety interactions suggest that selective pressures operate at very small spatial scales, with apple varieties often being planted within meters of each other and showing strong community differentiation. This close proximity of different varieties falls within yeast’s dispersal neighborhoods, as they are moved by fruit flies and wasps that commonly feed on, and move between, rotting apples. This scale-dependent community assembly has important implications for both theoretical understanding of microbial community composition and orchard management.

### H. uvarum Local (Mal)Adaptation

The complex patterns of *H. uvarum,* from strong local adaptation in some orchards to maladaptation in others, demonstrate the effects of rapid evolution at microgeographic scales. This aligns with previous findings of *H. uvarum* genetic differentiation in vineyards (Albertin et al. 2016), but this study extends this to even smaller geographic scales. Our results reveal multiple interesting patterns that require explanation: the variation among strains within populations, the contrasting patterns between orchards, and the interaction with apple varieties.

The strain-level variation in performance suggests that multiple adaptive solutions exist within populations. This intra-population variation might reflect temporal heterogeneity in selection pressures, as orchard conditions change seasonally and yearly, and yeast can evolve rapidly. Additionally, the existence of multiple apple varieties within each orchard could maintain genetic variation through habitat heterogeneity. The strong performance differences among strains (ranging from β=-0.106 to 0.198) indicate that there is substantial functional diversity within *H. uvarum*. These results align with those of Leducq et al (2014), who found that the ubiquitous yeast species *Saccharomyces paradoxus* contained abundant among-strain variation in fitness.

Maladaptation in some orchards might reflect several processes (Brady et al. 2019). Changes in management practices could create mismatches between historically adapted populations and current conditions. Another possibility is that apparent maladaptation could reflect adaptation to competing species within home orchards, but by measuring *H. uvarum* adaptation in monoculture, we only assessed its adaptation to the apple substrate, not to the complete environment including other yeast species. Strains that perform poorly in isolation might perform better in competitive community contexts. Additionally, since the yeast are actively degrading the apples, maladaptation (rotting the apple less) may be advantageous at times.

These results support theoretical predictions about rapid evolution’s importance in microbial systems while also revealing complexity in adaptation patterns. The variation in adaptive outcomes among orchards and varieties suggests that predicting evolutionary trajectories of yeast requires understanding the interaction between management practices, apple characteristics, species interactions, and respective abiotic factors of the orchards.

### Local adaptation impacts on community

The result of local adaptation driving community composition and not performance is a novel finding with important implications for eco-evolutionary dynamics. While *H. uvarum* local adaptation to the apple substrate strongly influenced which species co-occurred, particularly by suppressing *Pichia* species, it did not affect overall community performance. This pattern challenges predictions about local adaptation’s effects on communities.

Our results contrast with Pantel et al. (2015) and PedersenHart et al. (2019), who both found that rapid experimental evolution led to more even abundances in their respective communities. Similar to these studies, Gómez et al. (2016) found that experimental evolution of *Pseudomonas fluorescens* led to variable directional effects on other species’ abundances and also found that local adaptation of *P. fluroescens* increased overall culturable bacteria in compost. In contrast to all of these studies, we found that local adaptation made competitive hierarchies more extreme, resulting in *H. uvarum* dominating the community when locally adapted, while having no effect on overall community performance. However, in all of these contrasting cases, our study is different as we do not conduct evolution experimentally, and instead focus on characterizing the current phenotype of yeast that has been evolving to environmental conditions for hundreds of years. This could indicate that there could be temporal eco-evolutionary dynamics at play in these orchards that cannot be identified by a single snapshot in time.

The maintenance of community performance despite dramatic compositional changes supports the portfolio effect and the role of functional redundancy in communities (Schindler et al. 2015). We found that different community compositions, from those dominated by *H. uvarum* to more diverse assemblages, achieved similar functional outcomes regardless of *H. uvarum* local adaptation or maladaptation. This suggests there could be functional redundancy among yeast species in fruit degradation (Louca et al. 2018). However, the portfolio effect stresses the importance of diversity for buffering against environmental variation, and while those communities dominated by *H. uvarum* are functionally the same they may not be as robust to changes in the environment. There also is a possibility that there might be a maintenance of diversity through the meta-community of yeast in the air (Urban et al. 2008).

The monopolization hypothesis (Urban and De Meester 2009; De Meester et al. 2016) might help explain how *H. uvarum*’s adaptation leads to competitive dominance, but since we don’t test priority effects in this experiment the exact mechanism of their dominance is unclear. Our results, however, extend this framework by showing that monopolization doesn’t necessarily enhance or decrease community performance. In communities where *H. uvarum* was locally adapted, it could comprise up to 97.5% of the community while maintaining equivalent performance to far more diverse communities.

### Broader Implications

These findings enhance our understanding of eco-evolutionary dynamics while raising new questions about community assembly and performance. First, we demonstrate that local adaptation can drive community composition without affecting performance. The disconnect between composition and performance suggests we may need to reconsider how we predict community responses to environmental change (Reed and Martiny 2007).

Our results also demonstrate that microgeographic adaptation occurs even in highly mobile, rapidly evolving microbial species. Yeasts disperse via air currents, insects, and larger animals that visit multiple trees and orchards. Despite this high dispersal potential, which should theoretically homogenize populations especially within orchards, we observed significant local adaptation at the spatial scale of varieties within orchards, suggesting that strong selection pressures can overcome gene flow in this system and supporting work on microgeographic adaptation (Richardson et al. 2014). The complex patterns of adaptation and maladaptation across orchards suggest that evolutionary trajectories remain difficult to predict even in well-defined experimental systems with replicated clonal habitats. This has important implications for understanding how communities might respond to environmental change, because while species may adapt rapidly, their evolutionary trajectories may be complicated by a variety of interacting factors.

From an agricultural perspective, understanding how yeasts adapt to specific orchards and varieties could inform management practices. The strong effect of orchard-variety interactions suggests rotating fungicides might help manage yeast communities. However, the maintenance of community performance across different compositions suggests that even if a management strategy can eliminate many species, even those communities that are less diverse or not as adapted can rot apples equally as well.

Several questions emerge from this study. First, while we found strong effects of *H. uvarum* adaptation on community composition, we do not know the temporal dynamics of this process. Priority effects could alter these dynamics significantly, for example, would early colonization by maladapted strains lead to different community outcomes? Additionally, genomic analysis of *H. uvarum* strains could reveal whether similar genetic changes underlie parallel adaptation across orchards, providing insight into the predictability of evolution at microgeographic scales.

Future work should explore whether these patterns hold in other microbial systems and evaluate the impact of priority effects on these dynamics. Additionally, observing this community dynamic experimentally or collecting temporal data may elucidate the mechanisms by which we are seeing these results. While our results challenge some assumptions about diversity-stability relationships, they may be specific to systems with high functional redundancy and rapid adaptation. Testing these ideas in communities with more specialized species interactions could reveal important constraints on when local adaptation influences community function.

Our study demonstrates that adaptation of a dominant species can dramatically restructure community composition while community function remains robust. This decoupling of community composition from community performance that we observed has important implications for both evolutionary ecology and the evaluation of community responses to environmental change. It suggests predicting community responses to environmental change may not require large amounts of diversity, but rather an understanding of which species are present in a community and to what extent they have substitutable functions. In agricultural contexts, this implies that management practices targeting one or a few pest species may not necessarily alter functional outcomes if redundancy exists among community members. More broadly, our work highlights how rapid evolutionary processes operating at small spatial scales can influence biodiversity patterns without disrupting ecosystem functions, contributing to our understanding of the complex interplay between ecology and evolution in natural systems.

## Supplementary Figures and Tables

**Supplemental Figure 1:**
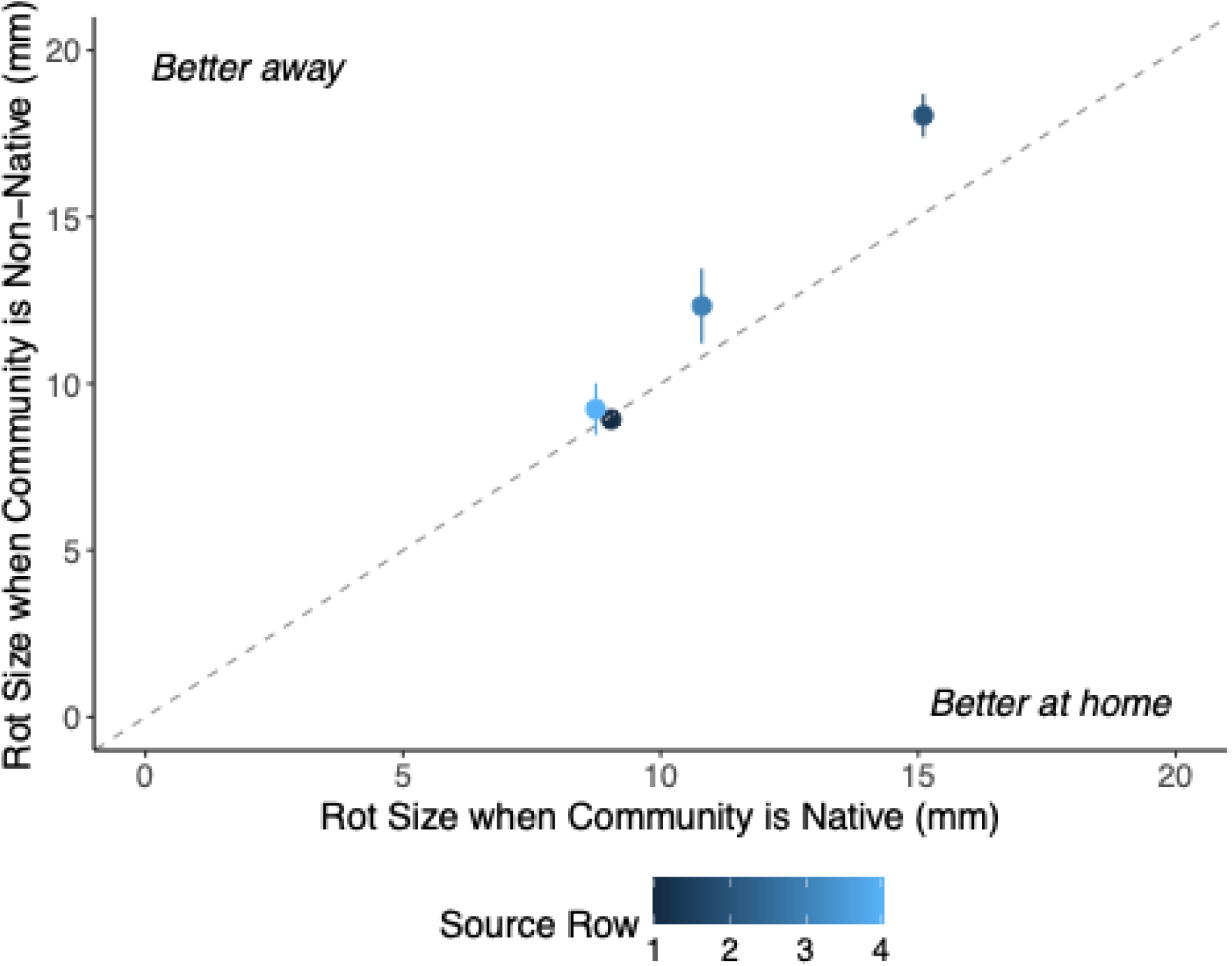
Community Performance in Native versus Non-native Row. Rot size produced by yeast communities in reciprocal transplant experiments by apple orchard row.

**Supplemental Figure 2:**
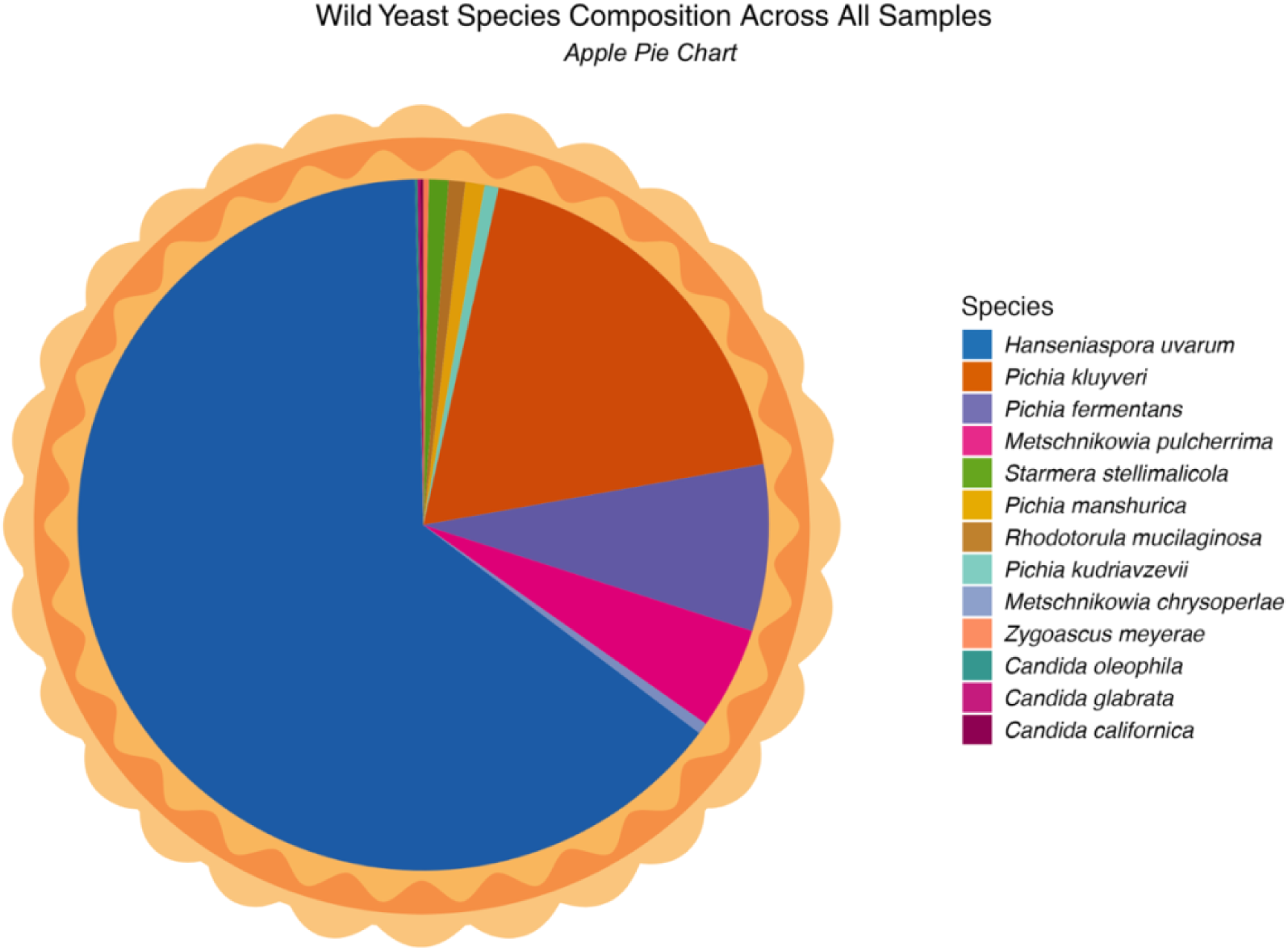
Apple Pie Chart of Wild Yeast Species Composition Across All Samples. Species composition across all orchards and varieties presented as a pie chart.

**Supplemental Table 1:**
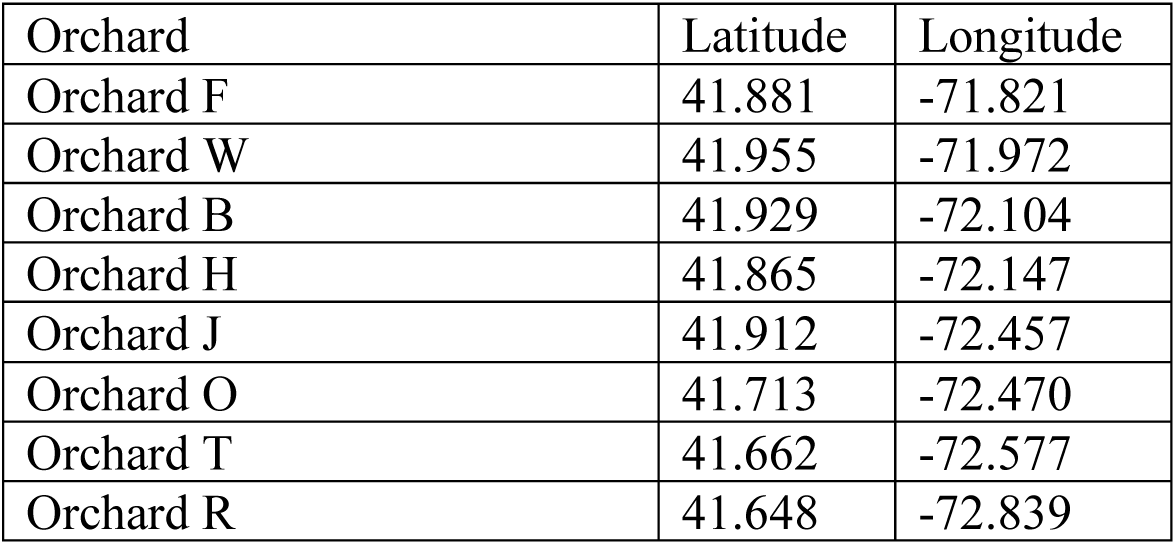
GPS Coordinates of the eight orchards evaluated in this study.

| Orchard | Latitude | Longitude |
| --- | --- | --- |
| Orchard F | 41.881 | -71.821 |
| Orchard W | 41.955 | -71.972 |
| Orchard B | 41.929 | -72.104 |
| Orchard H | 41.865 | -72.147 |
| Orchard J | 41.912 | -72.457 |
| Orchard O | 41.713 | -72.470 |
| Orchard T | 41.662 | -72.577 |
| Orchard R | 41.648 | -72.839 |

**Supplemental Table 2:** Community performance by tree row.

| a) community performance, home vs away (by yeast source) |  |  |  |
| --- | --- | --- | --- |
| **Factor** | **df** | ***F* value** | ***p* value** |
| Native/Non-native Row | 1 | 0.04 | 0.85 |
| Yeast Source Row | 3 | 0.08 | 0.78 |
| Native/Non-native Row* Source Row | 3 | 0.00 | 0.98 |
| b) community performance, local vs foreign (by apple source) |  |  |  |
| **Factor** | **df** | ***F* value** | ***p* value** |
| Native/Non-native Row | 1 | 0.01 | 0.92 |
| Apple Source Row | 3 | 0.08 | 0.78 |
| Native/Non-native Row* Source Row | 3 | 0.12 | 0.73 |

